# PlantiSMASH: automated identification, annotation and expression analysis of plant biosynthetic gene clusters

**DOI:** 10.1101/083535

**Authors:** Satria A. Kautsar, Hernando G. Suarez Duran, Kai Blin, Anne Osbourn, Marnix H. Medema

**Affiliations:** Bioinformatics Group, Wageningen University, 6708 PB Wageningen, The Netherlands; Teknik Informatika, Universitas Lampung, Jln. Sumantri Brojonegoro No. 01, Lampung 35141,Indonesia; The Novo Nordisk Foundation Center for Biosustainability, Technical University of Denmark, 2800 Kgs. Lyngby,Denmark; Department of Metabolic Biology, John Innes Centre, Norwich Research Park, Norwich, NR4 7UH,United Kingdom

## Abstract

Plant specialized metabolites are chemically highly diverse, play key roles in host-microbe interactions, have important nutritional value in crops and are frequently applied as medicines. It has recently become clear that plant biosynthetic pathway-encoding genes are sometimes densely clustered in specific genomic loci: biosynthetic gene clusters (BGCs). Here, we introduce plantiSMASH, a versatile online analysis platform that automates the identification of candidate plant BGCs. Moreover, it allows integration of transcriptomic data to prioritize candidate BGCs based on the coexpression patterns of predicted biosynthetic enzyme-coding genes, and facilitates comparative genomic analysis to study the evolutionary conservation of each cluster. Applied on 48 high-quality plant genomes, plantiSMASH identifies a rich diversity of candidate plant BGCs. These results will guide further experimental exploration of the nature and dynamics of gene clustering in plant metabolism. Moreover, spurred by the continuing decrease in costs of plant genome sequencing, they will allow genome mining technologies to be applied to plant natural product discovery.

The plantiSMASH web server, precalculated results and source code are freely available from http://plantismash.secondarymetabolites.org.

## INTRODUCTION

Across Planet Earth, bacteria, fungi and plants produce an immense diversity of specialized metabolites, each with their own specific ecological roles in the manifold interorganismal interactions in which they engage. This diverse specialized metabolism is a rich source of natural products that are used widely in medicine, agriculture and manufacturing. In bacteria and fungi, where genes for most specialized metabolic pathways are physically clustered in so-called biosynthetic gene clusters (BGCs), the rapid accumulation of genome sequences has revolutionized the process of natural product discovery: indeed, genome mining has now become a dominant method for the discovery of novel molecules (1–4). In the genome mining process, BGCs are computationally identified in genome sequences and then linked to compounds through functional analysis (e.g., using metabolomic data, chemical structure predictions, mutant libraries, and/or heterologous expression). Many sequence-based aspects of this genome mining procedure are facilitated by the online antiSMASH framework, which was launched in 2010 (5) and has seen continuous development since then (6, 7). The genome mining procedure has two main purposes: 1) finding biosynthetic genes for important known compounds to allow heterologous production through fermentation in industrial strains, and 2) identifying novel natural product chemistry guided by biosynthetic gene cluster diversity. Altogether, this development has appropriately been termed the ‘gene cluster revolution’ (1). In recent years, it has become clear that not only microbial, but also plant biosynthetic pathways are frequently chromosomally clustered: after the initial discoveries of the cyclic hydroxamic acid 2,4-dihydroxy-1,4-benzoxazin-3-one (DIBOA) and avenacin gene clusters (8, 9), around thirty plant BGCs have been discovered (10, 11). Together, they encode the production of a wide range of different compounds, including cyclic hydroxamic acids, di-and triterpenes, steroidal and benzylisoquinoline alkaloids, cyanogenic glucosides and polyketides. In the genome of the model plant species *Arabidopsis thaliana* alone, four BGCs have been linked to specific metabolites, and recent analyses based on epigenomic profiling indicate the presence of various additional uncharacterized ones (12). Various technological developments in eukaryote genome sequencing (13) are finally making complete plant genome sequencing feasible at larger scales: high-quality plant genome sequences for almost 100 species are already publicly available, and more or less complete genomes can be sequenced for as little as a 10-50k US dollars each. Hence, genome mining may become an important methodology in the study of plant natural products as well, and a realistic opportunity thus presents itself for the plant natural product research community to have a ‘gene cluster revolution’ of its own. Naturally, a key technology required to realize this is a computational framework specifically designed for the identification and analysis of plant BGCs. Importantly, tools available for bacterial and fungal genome mining do not suffice for plants (14), as 1) plant biosynthetic pathways involve unique enzyme families not found in bacteria and fungi; 2) not all plant biosynthetic pathways are clustered (e.g., anthocyanins (15)), so identification of a biosynthetic gene does not equal identification of a BGC; 3) intergenic distances in plant genomes are larger and much more variable (16–19); 4) plant genomes contain clustered groups of genes (e.g., tandem arrays) whose products do not constitute a pathway; 5) several plant pathways are split across more than one BGC (20, 21). Here, we introduce antiSMASH for plants (or ‘plantiSMASH’ in short), which has been designed to tackle each of these challenges. Through a comprehensive library of profile Hidden Markov Models (pHMMs) for enzyme families known to be involved in plant biosynthetic pathways, combined with CD-HIT clustering of predicted protein sequences belonging to the same family, it allows the efficient identification of genomic loci encoding multiple different (sub)families of specialized metabolic enzymes. Moreover, comparative genomic analysis as well as analysis of gene expression patterns within these candidate BGCs allow assessment of each locus for its likelihood to encode genes working together in one pathway. Finally, coexpression analysis between candidate BGCs and with other genes across the genome allows identification of biosynthetic pathways that are encoded on multiple loci. To exploit this new framework, we offer an initial analysis of BGC diversity across the plant kingdom, which showcases the presence of many complex biosynthetic loci in diverse species.

## MATERIAL AND METHODS

### A procedure for the identification of candidate plant biosynthetic gene clusters

The microbial version of antiSMASH (5) predicts BGCs by using HMMer (22) to identify specific (combinations of) signature protein domains that belong to scaffold-generating enzymes specific for a class of biosynthetic pathways. Subsequently, hit genes are used as anchors from which gene clusters are extended upstream and downstream by a specified extension distance.

Although very effective for detecting biosynthetic clusters on bacteria and fungi, this procedure is unfit to detect biosynthetic gene clusters in plants, for the reasons described above. To address these differences, a novel detection strategy was chosen (Figure 1): instead of identifying BGCs through the identification of core scaffold-generating genes alone, plantiSMASH identifies them by looking for all genes predicted to encode different types of biosynthetic enzymes, including those required for tailoring of the scaffold.

**Figure 1.**
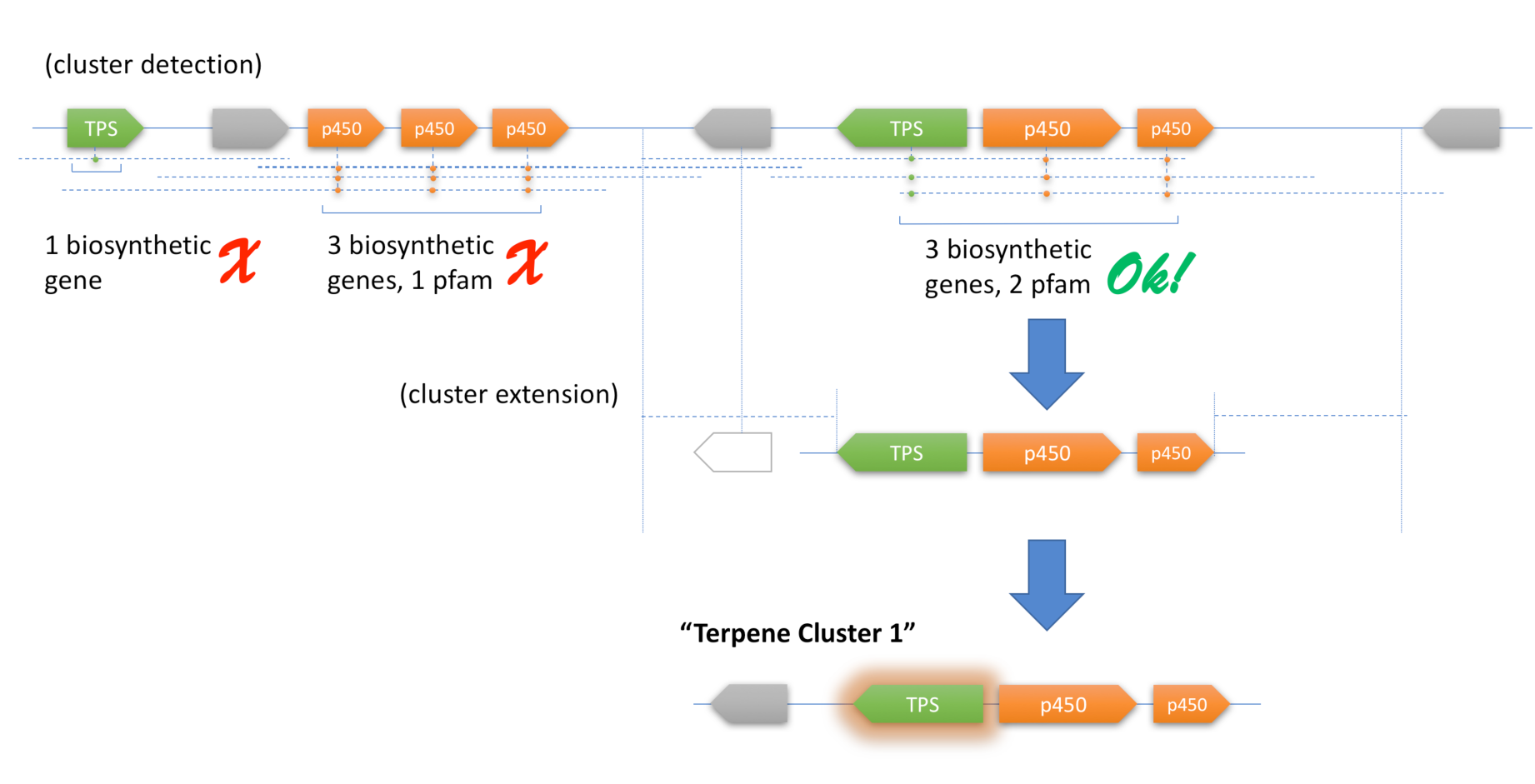
General strategy followed by plantiSMASH for the identification of plant BGCs. First, plantiSMASH will try to identify biosynthetic genes (having a hit on one of the 62 pHMMs) that are located in close proximity to each other. In particular, it will look for the co-occurrence of at least three biosynthetic enzyme-coding genes, comprising at least two different enzyme types. (Based on the results of the CD-HIT clustering of encoded protein sequences, closely related duplicate genes will only be counted once). Afterwards, identified cluster are extended to incorporate any flanking genes. Finally, each cluster is classified based on the presence of core enzymes (see SI Table 1). In this example, the detected cluster is assigned to the “Terpene” class due to the presence of a terpene synthase-encoding gene.

To determine what constitutes a high-potential candidate BGC, we make use of the recently proposed definition for plant BGCs as ‘genomic loci encoding genes for a minimum of three different types of biosynthetic reactions (i.e. genes encoding functionally different (sub)classes of enzymes)’ (14). (Albeit arbitrary, this definition correctly describes all known plant BGCs at the moment, and is open to improvement as more are discovered.) Accordingly, with default settings plantiSMASH defines clusters as loci where at least three different enzyme subclasses belonging to at least two different enzyme classes are co-located on the same locus. Enzyme classes are identified using profile Hidden Markov Models (pHMMs) specific for each class (SI Table 1); to count the number of subclasses of each enzyme class at a certain locus, the CD-HIT algorithm (23) is employed for sequence-based clustering to identify groups of sequences within an enzyme class with (by default) >50% mutual amino acid sequence identity. This successfully distinguishes potentially real BGCs from tandem repeat regions that are also frequently found in genomes (SI Table 2).

In order to identify all classes of biosynthetic enzymes known to be involved in plant specialized metabolic pathways, we performed a comprehensive literature search of previously characterized plant biosynthetic pathways, which resulted in a list of 62 protein domains that have been associated with specialized metabolic pathways in plants (see SI Table 1). Fifty-seven of these protein domains are represented by pHMMs from the Pfam database (24), and custom pHMMs were only generated for five enzyme families not (fully) covered by Pfam domains. We consciously refrained from attempting to construct custom pHMMs for all enzyme families known to be involved in plant biosynthetic pathways, as the limited amount of training data available would lead to an overly strict prediction system that would no longer be able to detect biosynthetic novelty; instead, we assume that the broad enzyme families covered by Pfam domains are likely to be biosynthetically involved if multiple enzymes from these different families are encoded together in the same locus. As in the microbial version of antiSMASH, the presence of genes predicted to encode signature enzymes (defined as enzymes that determine the chemical class of the end compound, such as terpene synthases) in a candidate BGC are used to assign a cluster to a biosynthetic class (see SI Table 3 for cluster rules). However, compared to the microbial version, the biosynthetic classes in ‘plantiSMASH’ are more of an approximation, since not all signature enzyme families used can be unequivocally used to predict the compound type; e.g., while strictosidine synthase (25) and norcoclaurine synthase (26) are well-characterized members of the Bet v1 enzyme family, it is not clear what proportion of this family have similar Pictet-Spenglerase(-like) catalytic activities.

Another particular challenge for BGC detection in plant genomes is the large variation in gene density that occurs not only between but also within plant genomes (16–19). Replacing the static kilobase distance cut-off of microbial antiSMASH by a fixed cut-off based on the maximum number of genes that lie between each pHMM hit also does not provide a solution, as BGCs would then be allowed to cross large repeat regions or even centromeres. Therefore, we chose an alternative more dynamic cut-off that is a linear function of local gene density (defined as the gene density of the ten genes nearest to a pHMM hit), and applies a multiplier to calculate the cut-off in kb that is optimal for that specific genomic region (see **SI Table 2 and SI Figures 1+2** for results illustrating calibration of the default).

### Flexible and user-friendly input and output

To obtain reliable BGC predictions, a high-quality annotation of gene features in a genome is essential. While we do make available the option to run GlimmerHMM (27) on plant genome sequences, performing de novo gene finding on a raw FASTA file is not desirable, given the relatively low accuracy of these procedures. Because, additionally, the GenBank and EMBL input formats previously accepted for antiSMASH are not available for many plant genomes, we now allow users to supply input also in FASTA+GFF3 format, currently the most widely used format for describing plant genome annotations. For this, we implemented a new module based on Biopython’s GFF parsing package (http://biopython.org/wiki/GFF_Parsing) capable of combining the CDS features from the sequence input sequence, if any, with those of a file compliant to the Generic Feature Format Version as defined by The Sequence Ontology in 2003 (https://github.com/The-Sequence-Ontology/Specifications/blob/master/gff3.md). To properly match GFF3 CDS features to their correct sequence, the module demands record names (chromosome/scaffold/contigs) to be identical in both inputs; the only exception being if both inputs only contain one record, in which case the requirement is instead that no feature has coordinates outside the sequence range. This new module allows plantiSMASH to be used with genomes that are only annotated with GFF3 files, such as many of those present in the Joint Genome Institute’s Phytozome database (28).

Based on the biosynthetic gene cluster predictions, a rich and interactive HTML output is generated (Figure 2), which is largely reminiscent of the output of microbial antiSMASH jobs (5). Additionally, genes in the visualization page for each candidate BGC are colored based on the class of enzymes encoded, and a legend is provided that details the color scheme. On mouse click, panels for each gene provide information on the pHMMs that have hits against it, as well as on the amino acid identity to homologous genes within the same locus as calculated by CD-HIT.

**Figure 2.**
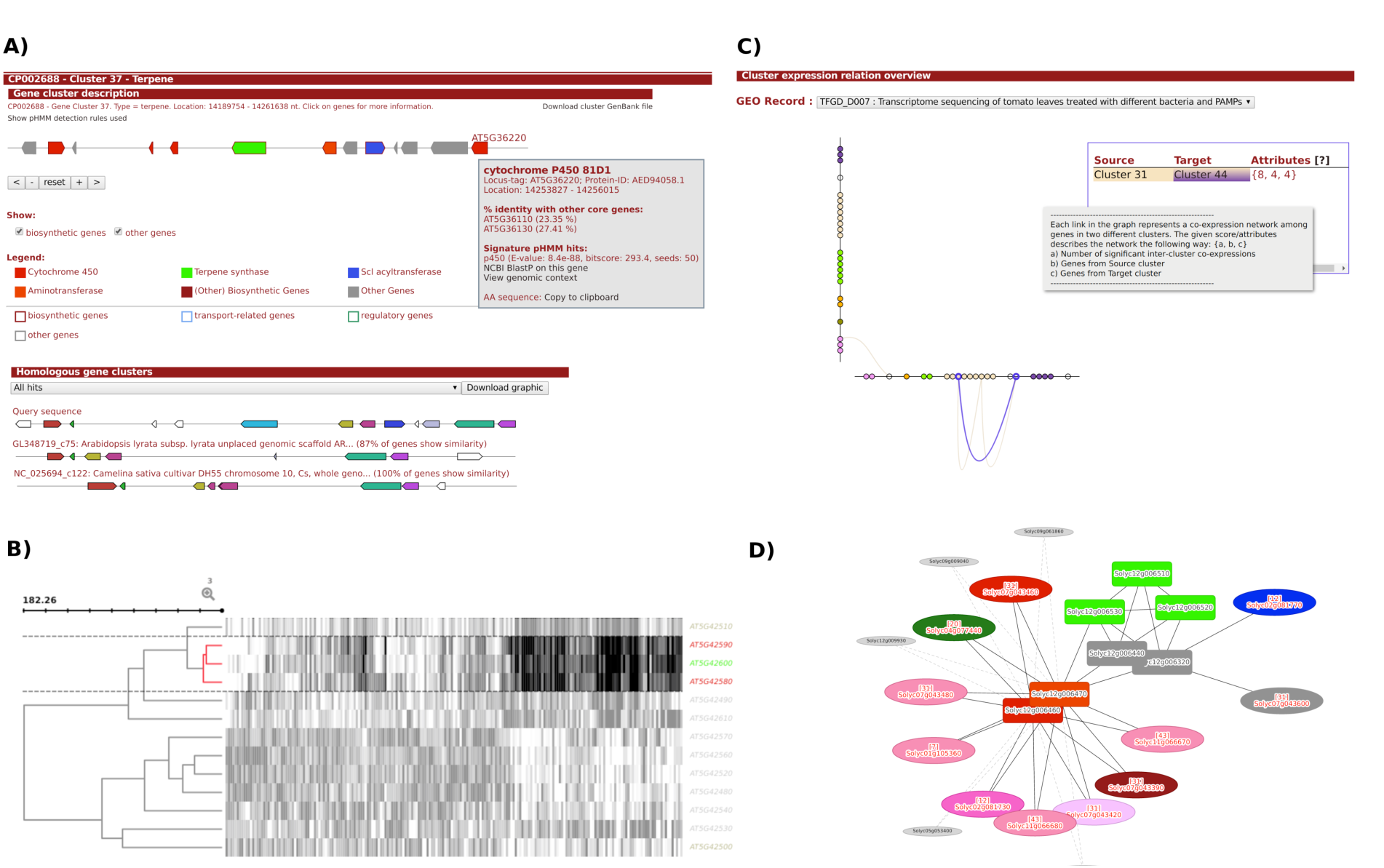
Outputs generated by the plantiSMASH pipeline. The figure illustrates several visualized outputs generated by plantiSMASH, as they appear for various biosynthetic gene clusters of known natural products. A) Visual overview generated for each gene cluster; in this case, the tirucalladienol cluster from *A. thaliana* (47) is shown. Gene annotations and pHMM hit details appear on mouse click. Also, ClusterBlast output showing alignment of homologous genomic loci across other genomes of related species is provided. B) Example of a gene expression heat map, showing coexpression among the core genes of the marneral BGC from *A. thaliana* (48) (and not with the flanking genes). C) Hive plot on the overview page, which highlights pairs of candidate BGCs which show many coexpression correlations between their genes; in this example view, the coexpression links between the two loci encoding α-tomatine biosynthesis in *Solanum lycopersicum* (20) are highlighted (clusters 31 & 44). D) Example ego network that summarizes coexpression correlations between members of the α-tomatine gene (cluster 44), as well as with genes in other gene clusters (including the other α-tomatine biosynthetic locus, cluster 31), and with genes elsewhere on the genome.

### Coexpression analysis identifies pathways within and between gene clusters

As plant scientists are just beginning to understand the phenomenon of metabolic gene clustering in plant genomes, it is currently unknown which proportion of genomic loci that encode multiple contiguous biosynthetic enzyme-encoding genes are bona fide BGCs in the sense that their constituent genes are involved in one specific pathway. One powerful strategy to predict whether genes are involved in the same pathway is the use of coexpression analysis, in which their expression patterns are compared across a wide range of samples. This strategy has proven very effective in the de novo identification of gene sets involved in biosynthetic pathways, even if they are not physically clustered on the chromosome (29).

To allow detailed investigation of whether genes in a cluster show coexpression, we added a dedicated analysis module: CoExpress. This module reads transcriptomic datasets, either in SOFT format (from the NCBI Gene Expression Omnibus) or in comma-separated (CSV) format, and generates powerful visualizations of these data for each candidate BGC. Because combining many datasets into one coexpression analysis may blot out coexpression signals that are very specific to certain biological or chemical treatments (which often highly specifically incite expression of plant specialized metabolic pathways), we designed the module in such a way that it visualizes one transcriptomic dataset at a time. This has the added value that the user can browse through multiple datasets and can individually assess specific samples that are linked to a treatment of interest. The visualizations of within-cluster coexpression patterns are twofold: First, a hierarchically clustered heatmap visualization, plotted using a modified version of the InCHlib (http://www.openscreen.cz/software/inchlib/home) JavaScript library, offers a direct view of patterns in and relationships between the supplied normalized gene expression values. The dendrogram is generated using a coexpression distance metric with a complete-linkage hierarchical clustering method. In this metric, the Pearson Correlation Coefficient (PCC) is transformed directly into a distance value scaled from 0 to 200 (0 for PCC = 1, or positively correlated, and 200 for PCC = −1, or negatively correlated). In order to make correlations maximally visible, the color scheme is normalized per gene (row) by default; however, the user can also select for the color scheme to be normalized by sample (column). Second, a gene cluster-specific coexpression network (30) (with a default distance based cutoff of < 50, dynamically adjustable) summarizes the correlations and helps to identify specific groups of genes in the locus that are highly coexpressed: these occur as connected components with high numbers of edges.

Coexpression analysis is not just useful for analysis of functional connections within a candidate BGC, but also allows prediction of functional links with other genomic loci. It is now well-understood that several plant BGCs do not act alone, but rather in concert with another BGC or with individual enzyme-coding genes elsewhere on the genome (11). Therefore, plantiSMASH leverages coexpression data to offer two analyses that identify these trans-genomic interactions: First, the BGC-specific coexpression network can be extended to display a first-order ego network that incorporates genes elsewhere on the genome that either 1) are members of another candidate BGC and show high gene expression correlation (> 0.9 PCC) with at least one gene in the BGC, or 2) contain a ‘biosynthetic’ domain (defined as being one of the domains in **SI Table 1**) and show high gene expression correlation with at least two genes in the BGC, at least one of which being a biosynthetic gene itself. Second, interactions between candidate BGCs are summarized in a hive plot, in which pairs of clusters are connected by an edge if the genes of both clusters create at least one subnetwork that satisfies the following criteria: 1) All nodes belong to the same Louvain community (31), as determined by analyzing the full coexpression network of all candidate clusters’ genes; 2) All nodes have a transitivity greater than zero; 3) The subnetwork contains at least two genes from each cluster; 4) The subnetwork contains at least one gene per cluster that has a biosynthetic domain; and 5) The subnetwork contains at least three genes with a biosynthetic domain. This highlights arrangements of pairs of clusters that may be linked functionally via coexpression, and is reminiscent of the characterized α-tomatine biosynthetic pathway in *S. lycopersicum*, which is encoded in two separate clusters that are highly coexpressed (20).

All in all, the coexpression analysis of candidate BGCs allows effective prioritization for, e.g., heterologous expression studies. Yet, it should still be kept in mind that loci that do not show high coexpression might still encode genes that are jointly involved in a biosynthetic pathway, e.g., if the transcriptomic samples available do not include any treatments that induce the expression of the pathway, or if expression of the pathway is sequestered either spatially across tissues or in terms of timing.

### Comparative genomic analysis shows conservation and diversification

Comparing a candidate BGC with homologous genomic loci in other plant genomes can give important information on its evolutionary conservation or diversification. Whereas strong conservation of clusteredness across larger periods of evolutionary time may point to a selective advantage of clustering for these genes, diversification of BGCs by co-option of other enzyme-coding genes may give clues to finding novel variants of natural products that have been generated through directional pathway evolution. In order to facilitate such comparative analysis on a case-by-case basis, we constructed a plant-specific version of the antiSMASH ClusterBlast module. To do so, we ran plantiSMASH on a collection of all publicly available plant genomes, obtained from NCBI’s GenBank, JGI’s Phytozome and Kazusa (32). In order to avoid cases where loci homologous to detected candidate BGCs would not be included in the database by not satisfying the identification criteria, the thresholds for this search were lowered to find all genomic loci with two or more different enzymes, where the CD-HIT cut-off was also set to a generously inclusive level of 0.9. A total of 7,978 genomic loci were thus included in the plant ClusterBlast database. As in the microbial version of antiSMASH, the translated protein sequence of each predicted gene in a candidate BGC is searched against this database using the DIAMOND algorithm (33), and genomic loci are sorted based on the number of hits, conserved synteny and cumulative bit score. To also facilitate direct comparison with known plant BGCs, all plant BGCs with known products for which the sequence was available were added to the MIBiG repository (34), which allows users to find similarities between newly identified and known clusters with the KnownClusterBlast module of antiSMASH.

### Precomputed results allow fast access to comprehensive plantiSMASH results

In order to allow users to directly access plantiSMASH results for publicly available plant genomes, runs for 47 high-quality plant genomes were precomputed and made available online at http://plantismash.secondarymetabolites.org (**SI Table 4**). Importantly, publicly available gene expression datasets with sufficient numbers of samples to be suitable for coexpression analysis were loaded into these results. In total, 73 transcriptomic datasets were included for five species: *Arabidopsis thaliana*, *Solanum lycopersicum*, *Oryza sativa*, *Zea mays* and *Glycine max* (**SI Tables 5-7**). Sequences that are not publicly available (as well as available sequences with custom transcriptomic datasets) can be analyzed directly using the plantiSMASH web server at http://plantismash.secondarymetabolites.org. In this way, plantiSMASH results for all kinds of genomes and transcriptomes are optimally available to users.

## RESULTS AND DISCUSSION

### PlantiSMASH successfully detects all experimentally characterized plant biosynthetic gene clusters

Even though only a relatively small set of plant BGCs has been discovered, these ~30 BGCs still present the best objective test case for the BGC detection algorithm. Importantly, they range from complex BGCs with many different enzyme-coding genes, such as the noscapine and cucurbitacin BGCs (21, 35), to relatively simple ones that only encode a couple of enzymes, such as the dhurrin and linamarin/lotaustralin BGCs (36). Of this set, only 19 BGCs have annotated sequence information publicly available. When plantiSMASH was run on a multi-GenBank file containing accurately annotated versions of these 19 known BGCs, all clusters were successfully detected with default settings. When run on different genome annotation versions available from GenBank or Phytozome, BGCs of low complexity (i.e., with a small number of enzyme-coding genes) were occasionally missed when key genes were missing from the structural annotations, or when many false positive gene assignments were present in the region of interest (affecting the dynamic gene density-based cut-off of plantiSMASH): for example, the linamarin BGC from *Lotus japonicus* was not detected in assembly/annotation version 3.0, while it was detected in the older version 2.5. This highlights the importance of using high-quality genome annotations supported by transcriptomic data when using plantiSMASH to search for BGCs of interest. Alternatively, the stand-alone version of plantiSMASH provides additional cut-off methods (e.g., raw distance-based or gene-count-based) that can be attempted as well to mitigate such issues.

### Plant genomes contain large numbers of complex biosynthetic gene clusters

When run on the 47 plant genomes for which chromosome-level assemblies are currently available on either NCBI or Phytozome, plantiSMASH found a wide variety of candidate BGC numbers across plant taxonomy (Figure 3). In general, the numbers of candidate BGCs were relatively even between monocots and dicots (while very low in the only moss genome included), while the largest numbers of BGCs were found in dicot genomes. These outliers all corresponded to recent (partial) genome amplification events, such as in the case of *Camelina sativa* (37) with 88 candidate BGCs, *Brassica napus* (38) with 68 candidate BGCs and *Glycine max* (39) with 76 candidate BGCs.

**Figure 3.**
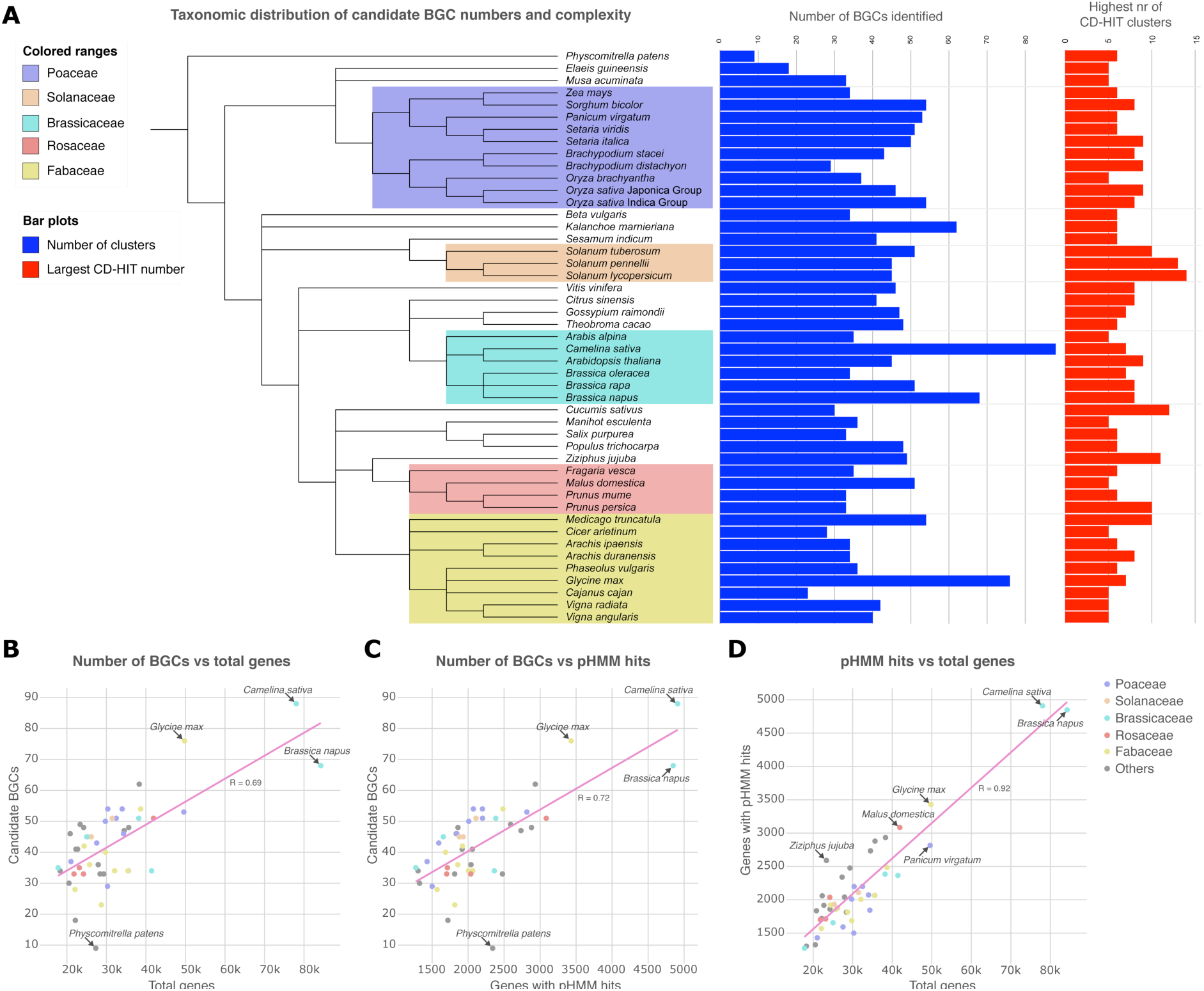
Numbers of candidate BGCs identified across the Plant Kingdom. A) PlantiSMASH BGC predictions plotted onto a phylogenetic tree of plant species for which chromosome-level genome assemblies are available. The blue bars indicate the number of candidate BGCs per genome, the red bars indicate the most complex candidate BGC identified in each species (in terms of the number of unique enzymes encoded, as defined by CD-HIT groups). B) Number of candidate BGCs plotted versus the total number of genes; as expected, more BGCs are found in larger genomes. Outliers represent genomes that have recently undergone whole-genome duplication, and the moss *Physcomitrella patens*, in the genome of which only a very low number of candidate BGCs is found. C) Number of candidate BGCs plotted versus the number of genes with pHMM hits to biosynthetic domains. D) Number of genes with biosynthetic domains plotted against the total number of genes; a linear correspondence is largely observed.

In many plant genomes, candidate BGCs of high complexity were identified, with as many as seven or eight different enzymatic classes encoded in the same tight genomic region. These constitutions are clearly non-random and make it promising to study candidate BGCs even in the absence of coexpression data. Dozens of such complex BGCs were found, which cover all known as well as putative pathway classes; examples are provided in Figure 4.

**Figure 4.**
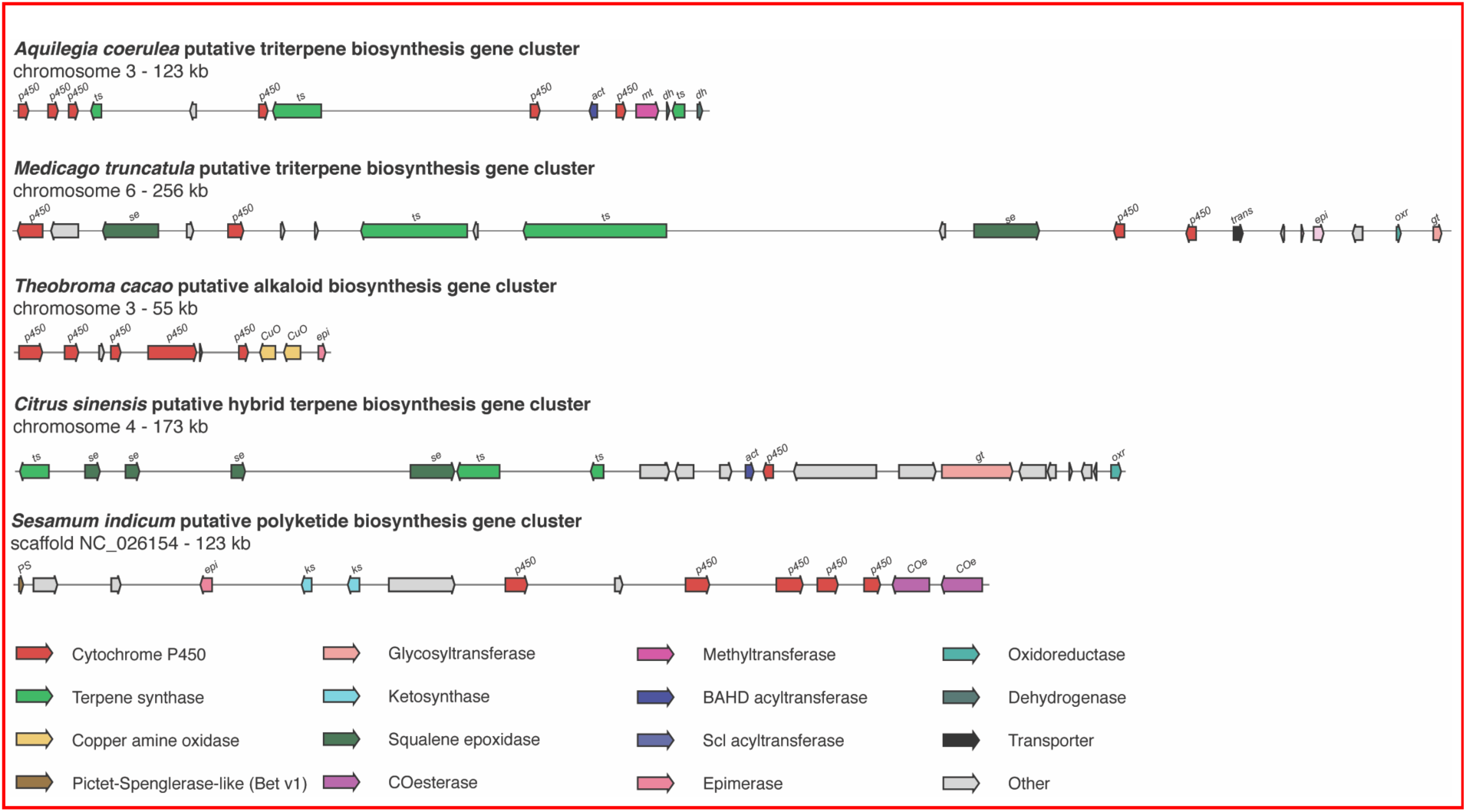
Example candidate BGCs identified by plantiSMASH. Five example candidate BGCs are shown, which cover a diverse range of enzymatic classes. Dozens of candidate BGCs of comparable complexity can be found across the precomputed plantiSMASH results that are available online.

### Coexpression patterns can guide BGC prioritization

We subjected the candidate BGCs identified in the genome of *A. thaliana* to a more detailed statistical analysis using within-cluster coexpression in a merged transcriptomic dataset. For this, we compiled two sets of gene expression datasets, one containing transcriptomic experiments of biological treatments (defense; **SI Table 5**) and one containing experiments of hormone treatments and non-biological stress inductions (**SI Table 6**). Together, these datasets comprise transcriptomic measurements of 1047 samples. The Mann–Whitney U one-sided test was selected to test which of the *A. thaliana* BGCs have a statistically greater within-cluster coexpression distribution than the genome’s background coexpression distribution. Given a BGC consisting of *x* genes, the background distribution for the statistical test of this cluster contains all PCCs between pairs of genes that are *x-1, x-2, …, 0* genes away from each other across the entire genome (except predicted BGCs). nly genes observed in all transcriptomic experiments were allowed in the test, and only PCCs between genes that each have a Median Absolute Deviation (MAD) larger than 0 are added to the distributions. Lastly, the CD-HIT algorithm was run on the entire *A. thaliana* proteome at 0.5 identity cutoff (same as plantiSMASH’s default) to cluster all similar enzymes. The same statistical tests were repeated afterwards, but this time discarding PCCs between genes that code for enzymes within the same CD-HIT cluster, ensuring both distributions only include coexpression of genes that produce enzymes of different classes, which more accurately resembles the type of interactions desired in a bona fide BGC. The results of these experiments (**SI Table 8** and SI Figure 3) show that a significance level of 0.05, 11 predicted BGCs showed statistically higher within-cluster coexpression than their respective background distribution even when discarding coexpression between genes in the same CD-HIT cluster. This list includes the four known A. thaliana BGCs, encoding the biosynthetic pathways for arabidiol/baruol (P=2.92e-40), thalianol (P=1.94e-17), marneral (P=7.03e-10) and tirucalla-7,24-dien-3β-ol (P=1.10e-4), which corroborates that coexpression is a valid criterion to prioritize functional BGCs.

There are several explanations for the fact that strong coexpression is observed for some candidate BGCs but not others. A first explanation is that their coordinated expression is induced by conditions not included in these transcriptomic experiments; in other words, absence of evidence of coexpression is not evidence of absence of coexpression. A second explanation is that a number of candidate BGCs probably do not encode entire consistently coexpressed biosynthetic pathways by themselves; evidence for this comes from an analysis of characterized enzyme-coding genes inside these candidate BGCs (**SI Table 9**); e.g., *AT1G24100* and *AT5G57220*, which occur in two different candidate BGCs, are known to be involved in two different branches of glucosinolate biosynthesis (40, 41), a complex multifurcated pathway that shows only partial and fragmented genomic clustering. Contrary to what might be expected, however, there was no strong correlation (R=0.004, and P=0.64 when fitting linear regression) of coexpression with cluster size, which suggests that the default plantiSMASH BGC prediction cut-offs are not set too inclusively.

All in all, coexpression analysis provides a powerful tool to prioritize the candidate BGCs detected by plantiSMASH that are most likely to encode functional pathways.

## CONCLUSIONS

The highly automated discovery of candidate BGCs by plantiSMASH and the powerful visualizations of coexpression data that allow their prioritization present a key technological step in the route towards high-throughput genome mining of plant natural products. As plant genome sequencing and assembly technologies continue to improve at a rapid pace, it is likely that high-quality plant genomes for thousands of species will soon be available; hence, ‘clustered’ biosynthetic pathways present low-hanging fruits for the discovery of novel molecules. Empowered by synthetic biology tools and powerful heterologous expression systems in yeast and tobacco (42–46), this will likely make it possible to scale up plant natural product discovery tremendously.

Continued development of the antiSMASH/plantiSMASH framework in the future is needed to further accelerate this process: e.g., the development of (machine-learning) algorithms that predict substrate specificities of key enzymes like terpene synthases, and the systematic construction of pHMMs for automated subclassification of complex enzyme families such as cytochrome P450s and glycosyltransferases, will allow more powerful predictions of the natural product structural diversity encoded in diverse BGCs. Additionally, detailed evolutionary genomic analysis of the phenomenon of gene clustering, including BGC birth, death and change processes, will further our understanding of how BGCs facilitate natural product diversification during evolution. As more plant BGCs are experimentally characterized, the algorithms will co-evolve with the knowledge gained, and more detailed class-specific cluster detection rules could be designed; moreover, it will become clearer what does and what does not constitute a bona fide BGC. Finally, when scientists further unravel the complexities of tissue-specific and differentially timed gene expression of plant biosynthetic pathways, we will learn more on how best to leverage coexpression data for biosynthetic pathway prediction. Thus, a more comprehensive understanding of the remarkable successes of evolution to generate an immense diversity of powerful bioactive molecules will hopefully make it possible for biological engineers to mimic nature’s strategies and deliver many useful new molecules for use in agricultural, cosmetic, dietary and clinical applications.

## ACKNOWLEDGEMENT

We thank Linh Nguyen for providing SI Table 9.

---

**SI Figure 1.**
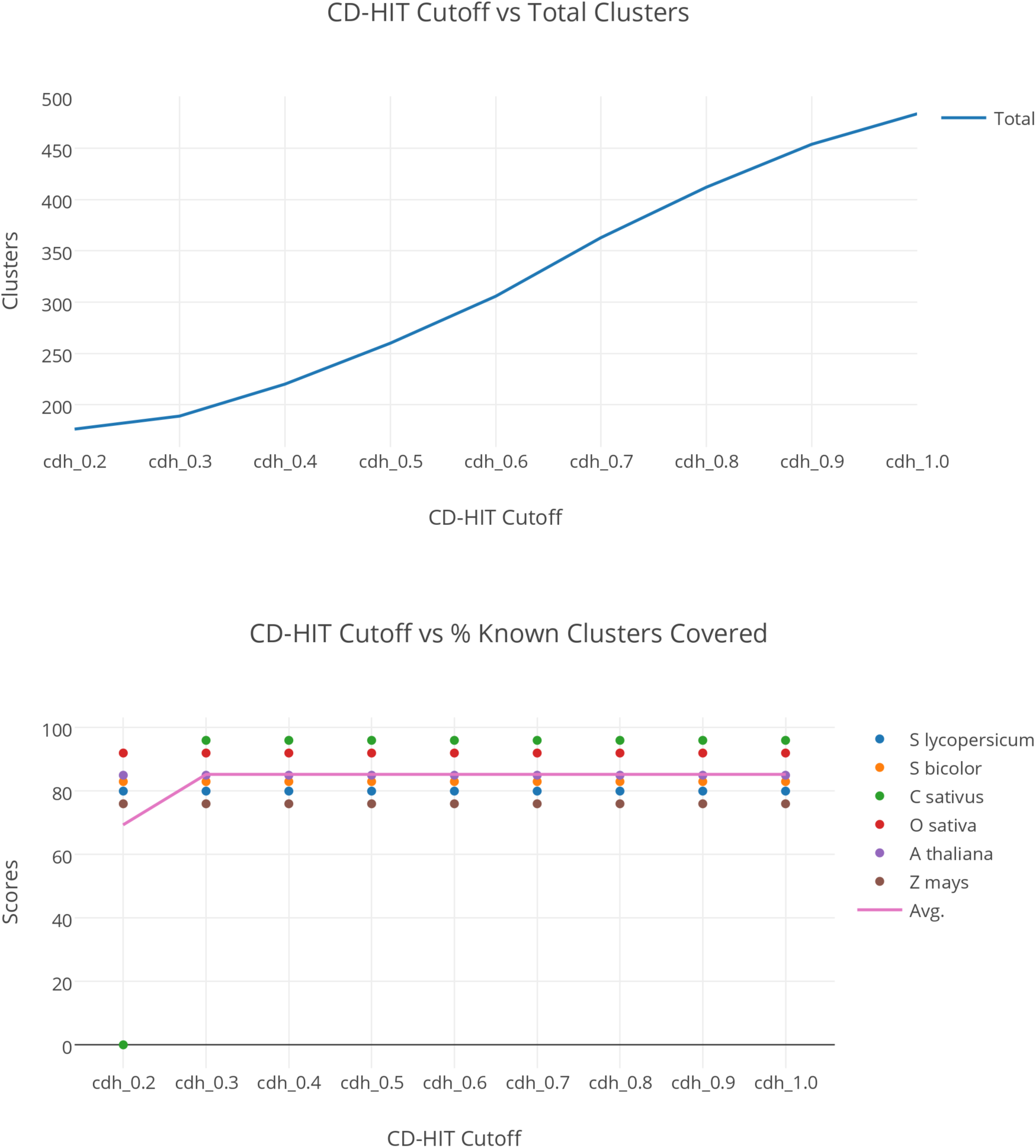
Effect of the CD-HIT cutoff parameter on the total number of predicted clusters and the coverage of known clusters. Coverage scores are calculated by comparing results of detected known clusters with literature (+1 for matching gene, −0.5 for absent gene, and -0.1 for extra gene; then converted to a percentage ratio). The default cut-off was chosen based on manual inspection of clusters that were gained/lost when changing the value of the parameter.

**SI Figure 2.**
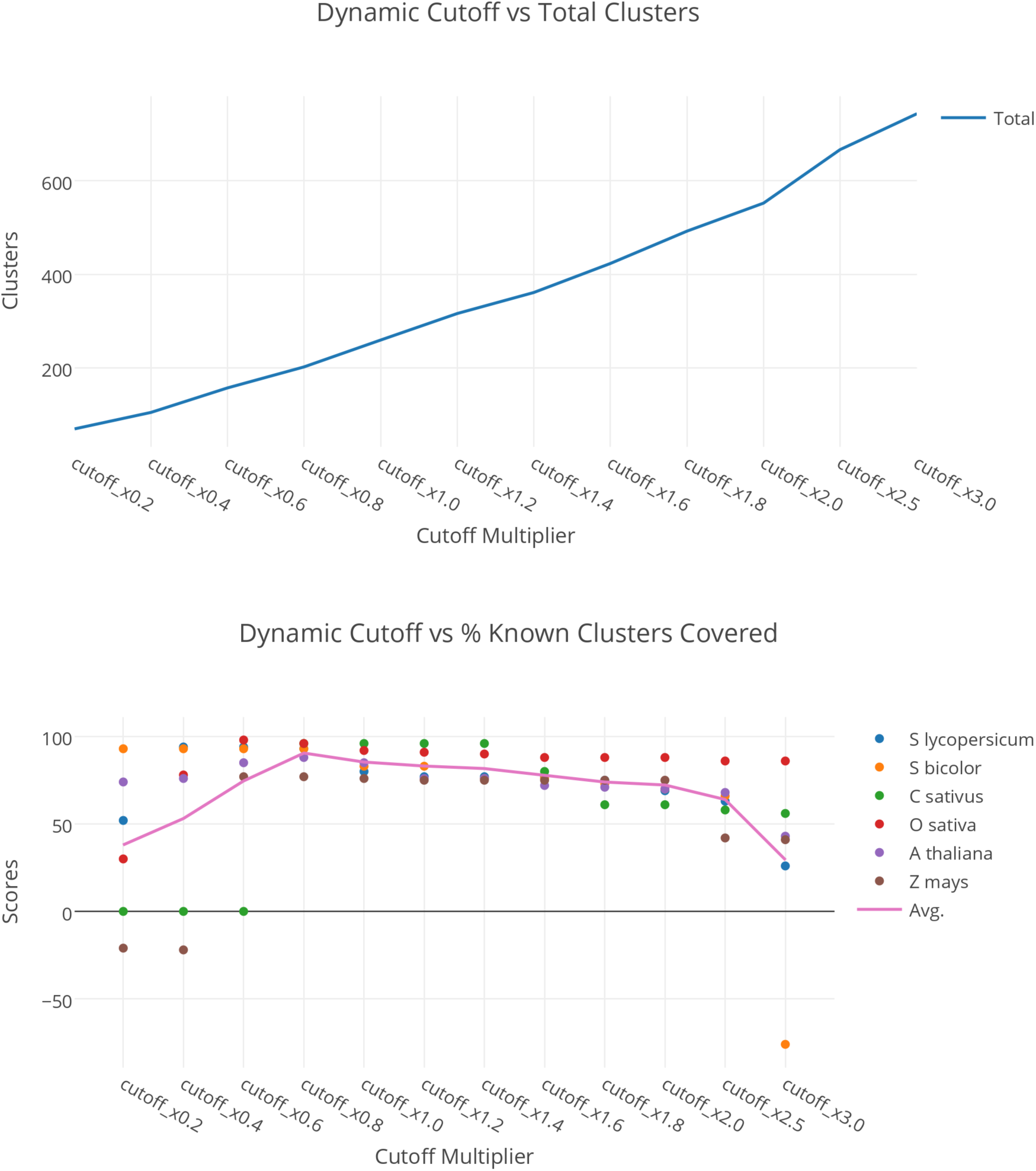
Effect of the dynamic cutoff parameter on the total number of predicted clusters and the coverage of known clusters. The default cut-off was chosen based on manual inspection of clusters that were gained/lost when changing the value of the parameter. Coverage scores are calculated by comparing results of detected known clusters with literature (+1 for matching gene, −0.5 for absent gene, and −0.1 for extra gene; then converted to a percentage ratio). The high score near the chosen default (lower panel) validates that the parameters effectively identify known gene clusters.

**SI Figure 3.**
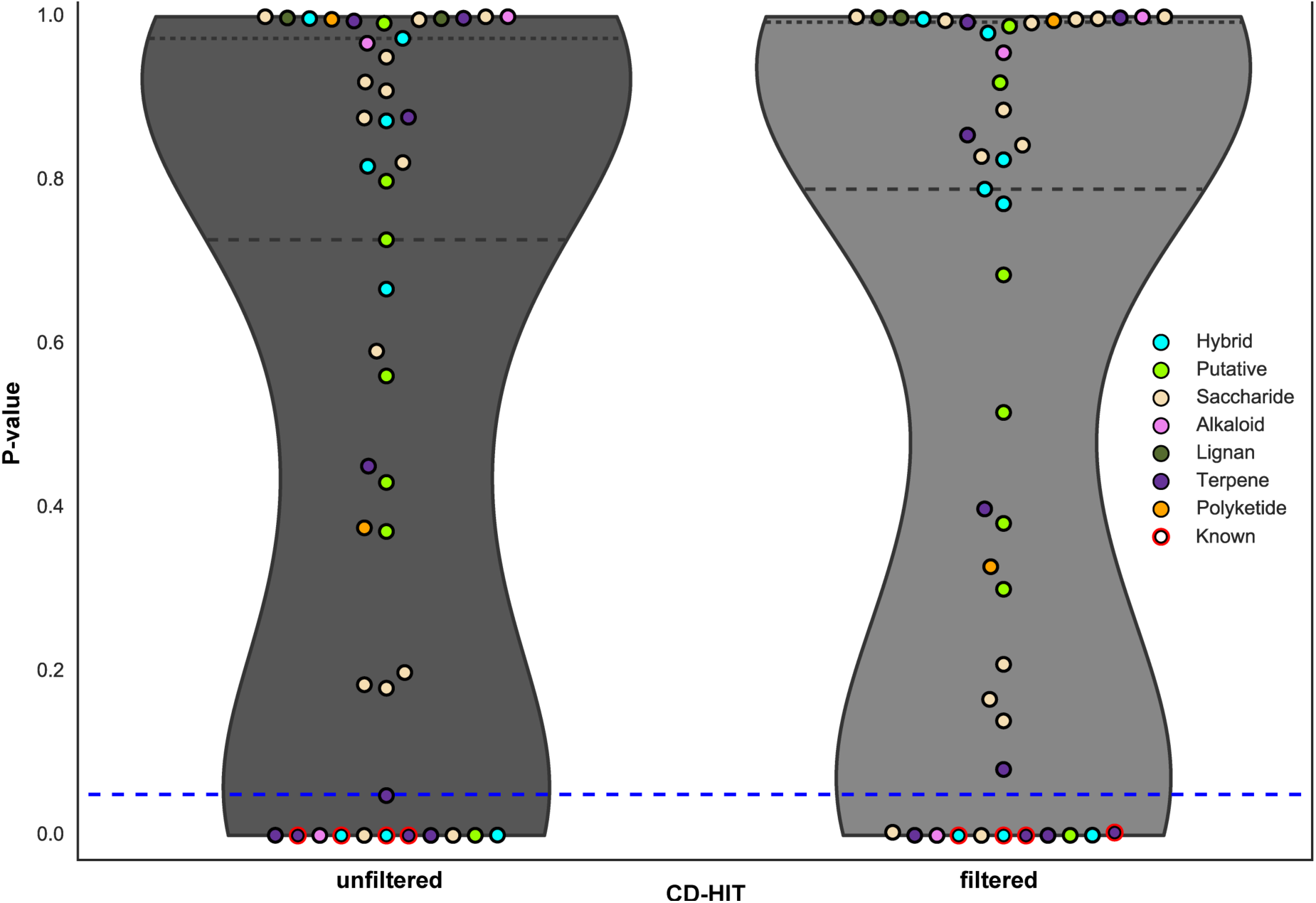
Mann-Whitney U test results of the PCC distribution of every *A. thaliana* cluster against a background distribution. Each circle represents a cluster, with fill colors denoting its assigned cluster type. Circles outlined in red denote known clusters (Marneral, thalianol, tirucalla and arabidiol/baruol). The blue dashed line indicates the significance level (P=0.05), and the black dashed lines the second and third quartile of each P distribution. Left: P distribution without discarding gene-pairs in the same CD-HIT cluster. Right: P distribution when discarding gene-pairs in the same CD-HIT cluster. Multiple testing correction was consciously refrained from given the fact that each comparison was among different samples, with each background distribution tailored to the cluster size; still, even applying a strict Bonferroni correction would only lower the number of significantly coexpressed clusters from 11 to 10.

## REFERENCES

1. Jensen, P.R. (2016) Natural Products and the Gene Cluster Revolution. Trends Microbiol. 24, 968–977.http://www.ncbi.nlm.nih.gov/pubmed/27491886 http://dx.doi.org/10.1016/j.tim.2016.07.006.

2. Medema, M.H. and Fischbach, M.A. (2015) Computational approaches to natural product discovery. Nat. Chem. Biol., 11, 639–648. http://www.ncbi.nlm.nih.gov/pubmed/26284671 http://dx.doi.org/10.1038/nchembio1884.

3. Rutledge, P.J. and Challis, G.L. (2015) Discovery of microbial natural products by activation of silent biosynthetic gene clusters. Nat. Rev. Microbiol., 13, 509–23. http://www.ncbi.nlm.nih.gov/pubmed/26119570 http://dx.doi.org/10.1038/nrmicro3496.

4. Ziemert, N., Alanjary, M. and Weber, T. (2016) The evolution of genome mining in microbes - a review. Nat. Prod. Rep., 33, 988–1005. http://www.ncbi.nlm.nih.gov/pubmed/27272205 http://dx.doi.org/10.1039/c6np00025h.

5. Medema, M.H., Blin, K., Cimermancic, P., de Jager, V., Zakrzewski, P., Fischbach, M.A., Weber, T., Takano, E. and Breitling, R. (2011) antiSMASH: rapid identification, annotation and analysis of secondary metabolite biosynthesis gene clusters in bacterial and fungal genome sequences. Nucleic Acids Res., 39, W339–W346. http://www.ncbi.nlm.nih.gov/pubmed/21672958http://dx.doi.org/10.1093/nar/gkr466.

6. Blin, K., Medema, M.H., Kazempour, D., Fischbach, M.A., Breitling, R., Takano, E. and Weber, T. (2013) antiSMASH 2.0—a versatile platform for genome mining of secondary metabolite producers. Nucleic Acids Res., 41, W204–W212. http://www.ncbi.nlm.nih.gov/pubmed/23737449 http://dx.doi.org/10.1093/nar/gkt449.

7. Weber, T., Blin, K., Duddela, S., Krug, D., Kim, H.U., Bruccoleri, R., Lee, S.Y., Fischbach, M.A., Muller, R., Wohlleben, W., et al. (2015) antiSMASH 3.0–a comprehensive resource for the genome mining of biosynthetic gene clusters. Nucleic Acids Res., 43, W237–W243.http://www.ncbi.nlm.nih.gov/pubmed/25948579 http://dx.doi.org/10.1093/nar/gkv437.

8. Frey, M., Chomet, P., Glawischnig, E., Stettner, C., Grün,S., Winklmair, A., Eisenreich, W., Bacher, A., Meeley, R.B., Briggs, S.P., et al. (1997) Analysis of a chemical plant defense mechanism in grasses. Science, 277, 696–9. http://www.ncbi.nlm.nih.gov/pubmed/9235894http://dx.doi.org/10.1126/science.277.5326.696.

9. Qi, X., Bakht, S., Leggett, M., Maxwell, C., Melton, R. and Osbourn, A. (2004) A gene cluster for secondary metabolism in oat: implications for the evolution of metabolic diversity in plants. Proc. Natl. Acad. Sci. U. S. A., 101, 8233–8. http://www.ncbi.nlm.nih.gov/pubmed/15148404http://dx.doi.org/10.1073/pnas.0401301101.

10. Nützmann,H.-W. and Osbourn, A. (2014) Gene clustering in plant specialized metabolism. Curr. Opin. Biotechnol., 26, 91–9. http://www.ncbi.nlm.nih.gov/pubmed/24679264http://dx.doi.org/10.1016/j.copbio.2013.10.009.

11. Nützmann,H.W., Huang, A. and Osbourn, A. (2016) Plant metabolic gene clusters - from genetics to genomics. New Phytol., 211, 771–789. http://www.ncbi.nlm.nih.gov/pubmed/27112429http://dx.doi.org/10.1111/nph.13981.

12. Yu, N., Nützmann,H.-W., MacDonald, J.T., Moore, B., Field, B., Berriri, S., Trick, M., Rosser, S.J., Kumar, S.V., Freemont, P.S., et al. (2016) Delineation of metabolic gene clusters in plant genomes by chromatin signatures. Nucleic Acids Res., 44, 2255–2265. http://www.ncbi.nlm.nih.gov/pubmed/26895889 http://dx.doi.org/10.1093/nar/gkw100.

13. VanBuren,R., Bryant, D., Edger, P.P., Tang, H., Burgess, D., Challabathula, D., Spittle, K., Hall, R., Gu, J., Lyons, E., et al. (2015) Single-molecule sequencing of the desiccation-tolerant grasshttp://www.ncbi.nlm.nih.gov/pubmed/26560029http://dx.doi.org/10.1038/nature15714.

14. Medema, M.H. and Osbourn, A. (2016) Computational genomic identification and functional reconstitution of plant natural product biosynthetic pathways. Nat. Prod. Rep., 33, 951–62. http://www.ncbi.nlm.nih.gov/pubmed/27321668http://dx.doi.org/10.1039/c6np00035e.

15. Shi, M.-Z. and Xie, D.-Y. (2014) Biosynthesis and metabolic engineering of anthocyanins in Arabidopsis thaliana. Recent Pat. Biotechnol., 8, 47–60.http://www.ncbi.nlm.nih.gov/pubmed/24354533http://dx.doi.org/10.2174/1872208307666131218123538.

16. Ibarra-Laclette,E., Lyons, E., Hernández-Guzmán,G., Pérez-Torres,C.A., Carretero-Paulet,L., Chang, T.-H., Lan, T., Welch, A.J., Juárez,M.J.A., Simpson, J., et al. (2013) Architecture and evolution of a minute plant genome. Nature, 498, 94–8http://www.ncbi.nlm.nih.gov/pubmed/23665961http://dx.doi.org/10.1038/nature12132.

17. Keller, B. and Feuillet, C. (2000) Colinearity and gene density in grass genomes. Trends Plant Sci., 5, 246–51. http://www.ncbi.nlm.nih.gov/pubmed/10838615http://dx.doi.org/10.1016/S1360-1385(00)01629-0.

18. Kellogg, E.A. and Bennetzen, J.L. (2004) The evolution of nuclear genome structure in seed plants. Am. J. Bot., 91, 1709–25. http://www.ncbi.nlm.nih.gov/pubmed/21652319http://dx.doi.org/10.3732/ajb.91.10.1709.

19. Sandhu, D. and Gill, K.S. (2002) Gene-containing regions of wheat and the other grass genomes. Plant Physiol., 128, 803–11. http://www.ncbi.nlm.nih.gov/pubmed/11891237http://dx.doi.org/10.1104/pp.010745.

20. Itkin, M., Heinig, U., Tzfadia, O., Bhide, A.J., Shinde, B., Cardenas, P.D., Bocobza, S.E., Unger, T., Malitsky, S., Finkers, R., et al. (2013) Biosynthesis of antinutritional alkaloids in solanaceous crops is mediated by clustered genes. Science, 341, 175–9. http://www.ncbi.nlm.nih.gov/pubmed/23788733http://dx.doi.org/10.1126/science.1240230.

21. Shang, Y., Ma, Y., Zhou, Y., Zhang, H., Duan, L., Chen, H., Zeng, J., Zhou, Q., Wang, S., Gu, W., et al. (2014) Plant science. Biosynthesis, regulation, and domestication of bitterness in cucumber. Science, 346, 1084–8. http://www.ncbi.nlm.nih.gov/pubmed/25430763http://dx.doi.org/10.1126/science.1259215.

22. Eddy, S.R. (2011) Accelerated Profile HMM Searches. PLoS Comput. Biol., 7, e1002195. http://www.ncbi.nlm.nih.gov/pubmed/22039361http://dx.doi.org/10.1371/journal.pcbi.1002195.

23. Fu, L., Niu, B., Zhu, Z., Wu, S. and Li, W. (2012) CD-HIT: accelerated for clustering the next-generation sequencing data. Bioinformatics, 28, 3150–2. http://www.ncbi.nlm.nih.gov/pubmed/23060610http://dx.doi.org/10.1093/bioinformatics/bts565.

24. Finn, R.D., Bateman, A., Clements, J., Coggill, P., Eberhardt, R.Y., Eddy, S.R., Heger, A., Hetherington, K., Holm, L., Mistry, J., et al. (2014) Pfam: the protein families database. Nucleic Acids Res., 42, D222–30.http://www.ncbi.nlm.nih.gov/pubmed/24288371 http://dx.doi.org/10.1093/nar/gkt1223.

25. Wu, F., Zhu, H., Sun, L., Rajendran, C., Wang, M., Ren, X., Panjikar, S., Cherkasov, A., Zou, H. and Stöckigt,J.(2012) Scaffold tailoring by a newly detected Pictet-Spenglerase activity of strictosidine synthase: from the common tryptoline skeleton to the rare piperazino-indole framework. J. Am. Chem. Soc., 134, 1498–500.http://www.ncbi.nlm.nih.gov/pubmed/22229634 http://dx.doi.org/10.1021/ja211524d.

26. Lee, E.-J. and Facchini, P. (2010) Norcoclaurine synthase is a member of the pathogenesis-related 10/Bet v1 protein family. Plant Cell, 22, 3489–503. http://www.ncbi.nlm.nih.gov/pubmed/21037103 http://dx.doi.org/10.1105/tpc.110.077958.

27. Majoros, W.H., Pertea, M. and Salzberg, S.L. (2004) TigrScan and GlimmerHMM: two open source ab initio eukaryotic gene-finders. Bioinformatics, 20, 2878–2879. http://www.ncbi.nlm.nih.gov/pubmed/15145805 http://dx.doi.org/10.1093/bioinformatics/bth315.

28. Goodstein, D.M., Shu, S., Howson, R., Neupane, R., Hayes, R.D., Fazo, J., Mitros, T., Dirks, W., Hellsten, U., Putnam, N., et al. (2012) Phytozome: a comparative platform for green plant genomics. Nucleic Acids Res., 40, D1178–86. http://www.ncbi.nlm.nih.gov/pubmed/22110026 http://dx.doi.org/10.1093/nar/gkr944.

29. Rajniak, J., Barco, B., Clay, N.K. and Sattely, E.S. (2015) A new cyanogenic metabolite in Arabidopsis required for inducible pathogen defence. Nature, 525, 376–379. http://www.ncbi.nlm.nih.gov/pubmed/26352477 http://dx.doi.org/10.1038/nature14907.

30. Serin, E.A.R., Nijveen, H., Hilhorst, H.W.M. and Ligterink, W. (2016) Learning from Co-expression Networks: Possibilities and Challenges. Front. Plant Sci., 7, 444. http://www.ncbi.nlm.nih.gov/pubmed/27092161 http://dx.doi.org/10.3389/fpls.2016.00444.

31. Blondel, V., Guillaume, J., Lambiotte, R. and Lefebvre, E. (2008) Fast unfolding of communities in large networks. J Stat Mech, 10, P10008. http://dx.doi.org/10.1088/1742-5468/2008/10/P10008.

32. Sato, S., Nakamura, Y., Kaneko, T., Asamizu, E., Kato, T., Nakao, M., Sasamoto, S., Watanabe, A., Ono, A., Kawashima, K., et al. (2008) Genome Structure of the Legume, Lotus japonicus. DNARes., 15, 227–239. http://www.ncbi.nlm.nih.gov/pubmed/18511435 http://dx.doi.org/10.1093/dnares/dsn008.

33. Buchfink, B., Xie, C. and Huson, D.H. (2014) Fast and sensitive protein alignment using DIAMOND. Nat. Methods, 12, 59–60. http://www.ncbi.nlm.nih.gov/pubmed/25402007 http://dx.doi.org/10.1038/nmeth.3176.

34. Medema, M.H., Kottmann, R., Yilmaz, P., Cummings, M., Biggins, J.B., Blin, K., De Bruijn, I., Chooi, Y.H., Claesen, J., Coates, R.C., et al. (2015) Minimum Information about a Biosynthetic http://www.ncbi.nlm.nih.gov/pubmed/26284661 http://dx.doi.org/10.1038/nchembio.1890

35. Winzer, T., Gazda, V., He, Z., Kaminski, F., Kern, M., Larson, T.R., Li, Y., Meade, F., Teodor, R., Vaistij, F.E., et al. (2012) A Papaver somniferum 10-gene cluster for synthesis of the anticancer alkaloid noscapine. Science, 336, 1704–8. http://www.ncbi.nlm.nih.gov/pubmed/22653730 http://dx.doi.org/10.1126/science.1220757.

36. Takos, A.M., Knudsen, C., Lai, D., Kannangara, R., Mikkelsen, L., Motawia, M.S., Olsen, C.E., Sato, S., Tabata, S., Jørgensen,K., et al. (2011) Genomic clustering of cyanogenic glucoside biosynthetic genes aids their identification in Lotus japonicus and suggests the repeated evolution of this chemical defence pathway. Plant J., 68, 273–86. http://www.ncbi.nlm.nih.gov/pubmed/21707799 http://dx.doi.org/10.1111/j.1365-313X.2011.04685.x.

37. Kagale, S., Koh, C., Nixon, J., Bollina, V., Clarke, W.E., Tuteja, R., Spillane, C., Robinson, S.J., Links, M.G., Clarke, C., et al. (2014) The emerging biofuel crop Camelina sativa retains a highly undifferentiated hexaploid genome structure. Nat. Commun., 5, 3706. http://www.ncbi.nlm.nih.gov/pubmed/24759634 http://dx.doi.org/10.1038/ncomms4706.

38. Chalhoub, B., Denoeud, F., Liu, S., Parkin, I.A.P., Tang, H., Wang, X., Chiquet, J., Belcram, H., Tong, C., Samans, B., et al. (2014) Plant genetics. Early allopolyploid evolution in the post-Neolithic Brassica napus oilseed genome. Science, 345, 950–3. http://www.ncbi.nlm.nih.gov/pubmed/25146293 http://dx.doi.org/10.1126/science.1253435.

39. Schmutz, J., Cannon, S.B., Schlueter, J., Ma, J., Mitros, T., Nelson, W., Hyten, D.L., Song, Q., Thelen, J.J., Cheng, J., et al. (2010) Genome sequence of the palaeopolyploid soybean. Nature, 463, 178–83. http://www.ncbi.nlm.nih.gov/pubmed/20075913 http://dx.doi.org/10.1038/nature08670.

40. Grubb, C.D., Zipp, B.J., Ludwig-Müller,J., Masuno, M.N., Molinski, T.F. and Abel, S. (2004) Arabidopsis glucosyltransferase UGT74B1 functions in glucosinolate biosynthesis and auxin homeostasis. Plant J., 40, 893–908. http://www.ncbi.nlm.nih.gov/pubmed/15584955 http://dx.doi.org/10.1111/j.1365-313X.2004.02261.x.

41. Pfalz, M., Vogel, H. and Kroymann, J. (2009) The gene controlling the indole glucosinolate modifier1 quantitative trait locus alters indole glucosinolate structures and aphid resistance in Arabidopsis. Plant Cell, 21, 985–99. http://www.ncbi.nlm.nih.gov/pubmed/19293369 http://dx.doi.org/10.1105/tpc.108.063115.

42. Casini, A., Storch, M., Baldwin, G.S. and Ellis, T. (2015) Bricks and blueprints: methods and standards for DNA assembly. Nat. Rev. Mol. Cell Biol., 16, 568–76. http://www.ncbi.nlm.nih.gov/pubmed/26081612 http://dx.doi.org/10.1038/nrm4014.

43. Liu, W., Yuan, J.S. and Stewart, C.N. (2013) Advanced genetic tools for plant biotechnology. Nat. Rev. Genet., 14, 781–793. http://www.ncbi.nlm.nih.gov/pubmed/24105275 http://dx.doi.org/10.1038/nrg3583.

44. Patron, N.J. (2014) DNA assembly for plant biology: techniques and tools. Curr. Opin. Plant Biol., 19, 14–9. http://www.ncbi.nlm.nih.gov/pubmed/24632010 http://dx.doi.org/10.1016/j.pbi.2014.02.004.

45. Patron, N.J., Orzaez, D., Marillonnet, S., Warzecha, H., Matthewman, C., Youles, M., Raitskin, O., Leveau, A., Farré,G., Rogers, C., et al. (2015) Standards for plant synthetic biology: a common syntax for exchange of DNA parts. New Phytol., 208, 13–9. http://www.ncbi.nlm.nih.gov/pubmed/26171760 http://dx.doi.org/10.1111/nph.13532.

46. Thimmappa, R., Geisler, K., Louveau, T., O’Maille,P. and Osbourn, A. (2014) Triterpene biosynthesis in plants. Annu. Rev. Plant Biol., 65, 225–57. http://www.ncbi.nlm.nih.gov/pubmed/24498976 http://dx.doi.org/10.1146/annurev-arplant-050312-120229.

47. Boutanaev, A.M., Moses, T., Zi, J., Nelson, D.R., Mugford, S.T., Peters, R.J. and Osbourn, A. (2014) Investigation of terpene diversification across multiple sequenced plant genomes. Proc. Natl. http://www.ncbi.nlm.nih.gov/pubmed/25502595 http://dx.doi.org/10.1073/pnas.1419547112.

48. Field, B. and Osbourn, A.E. (2008) Metabolic diversification––independent assembly of operon-like gene clusters in different plants. Science, 320, 543–547. http://www.ncbi.nlm.nih.gov/pubmed/18356490 http://dx.doi.org/10.1126/science.1154990.

